# Genome rearrangements and megaplasmid loss in the filamentous bacterium *Kitasatospora viridifaciens* are associated with protoplast formation and regeneration

**DOI:** 10.1101/629089

**Authors:** Karina Ramijan, Zheren Zhang, Gilles P. van Wezel, D. Claessen

## Abstract

Filamentous Actinobacteria are multicellular bacteria with linear replicons. *Kitasatospora viridifaciens* DSM 40239 contains a linear 7.8 Mb chromosome and an autonomously replicating plasmid KVP1 of 1.7 Mb. Here we show that lysozyme-induced protoplast formation of the multinucleated mycelium of *K. viridifaciens* drives morphological diversity. Characterization and sequencing of an individual revertant colony that had lost the ability to differentiate revealed that the strain had not only lost most of KVP1 but also carried lesions in the right arm of the chromosome. Strikingly, the lesion sites were preceded by insertion sequence elements, suggesting that the rearrangements may have been caused by replicative transposition and homologous recombination between both replicons. These data indicate that protoplast formation is a stressful process that can lead to profound genetic changes.

**Repositories:** Genomic sequence data for strain B3.1 has been deposited in the NCBI SRA database under accession code SAMN11514356.

## Introduction

Filamentous Actinobacteria are prolific producers of bioactive compounds. These metabolites are mostly used as weapons that provide protection against other microorganisms and phages in the environment [1–3]. This is particularly useful for filamentous organisms, given that they generally lack the ability to make flagella for escaping dangerous situations. In addition, these bacteria are able to generate resistant spores that can invade new environments after their dispersal. Germination of spores leads to the formation of 1-2 germ tubes, which grow by tip extension, thereby establishing filamentous cells called hyphae. Branching of hyphae leads to the formation of a multinucleated vegetative mycelium, which forages and acquires nutrients by decomposing polymeric substances. Stressful conditions (such as nutrient depletion) induce program cell death (PCD) of the mycelium, which in turn triggers morphological and chemical differentiation [4]. This developmental transition leads to the formation of specialized hyphae that grow into the air, and the onset of production of a suite of bioactive compounds [5]. Eventually, the aerial hyphae metamorphose into chains of grey-pigmented spores. Mutants that are unable to establish an aerial mycelium are called bald (*bld*), while those that are not capable to form spores are called white (*whi*) after their whitish color [6, 7].

Genome mining has been instrumental for the revival of drug discovery [8, 9]. Many of the biosynthetic gene clusters that specify bioactive natural products are contained on giant linear plasmids [10–13]. Although linear replicons are rare in many bacterial taxa, they are common in Actinobacteria [14, 15]. In fact, *Streptomyces* chromosomes (between 8 and 10 Mb in size) are also linear and typically comprise a “core region” containing the essential genes, and two variable “arms” with lengths ranging from 1.5 Mb to 2.3 Mb [16]. Like linear plasmids, the linear chromosomes are capped by terminal proteins bound to the 5’ end of the DNA [17]. The chromosomal ends are genetically unstable, and readily undergo large (up to 2 Mb) DNA rearrangements. Such rearrangements can lead to circularization of the chromosome, exchange of chromosomal arms or the formation of hybrid chromosomes due to recombination between the linear plasmids and the chromosome [18]. This wide range of genomic rearrangements is believed to be caused by transposition or homologous recombination, occurring actively within the chromosome or between the chromosome and linear plasmids [19]. Not surprisingly, these changes have profound effects on differentiation and specialized metabolite production [20, 21].

Here we characterized genetic instability in *Kitasatospora viridifaciens.* This tetracycline producer was originally classified within the genus *Streptomyces*, it was recently shown to belong to the genus *Kitasatospora* [22]. Protoplast formation and regeneration leads to the emergence of colonies that are no longer able to differentiate, which we attribute to the deletion of a 1.5 Mb segment of the right chromosomal arm and concomitant loss of most of the sequences contained on the large megaplasmid KVP1.

## Methods

### Strains and media

The strains used in this study (Table 1) are derivatives of *K. viridifaciens* DSM40239 (DSMZ). For protoplast preparation a spore suspension (10^6^ spores⋅ml^−1^) was grown for 48 hours in a mixture of TSBS-YEME (1:1 v/v) supplemented with 5 mM MgCl_2_ and 0.5% glycine. Protoplasts were prepared as described [23], with the difference that the lysozyme concentration was increased to 10 mg⋅ml^−1^. Serial dilutions of protoplasts were plated on R5 [23] or MYM medium [24] at 30°C. Regenerated protoplasts were streaked twice to single colonies on MYM before selecting the three independent bald colonies (B3.1, B3.2, and B3.3) that were further analyzed. Bald colonies were used as inoculum on liquid cultures of TSBS. Genomic DNA was isolated after two days of growth at 30°C.

**Table 1.** Strains used in this study

| Strains | Characteristics | Genotype | Reference |
| --- | --- | --- | --- |
| DSM 40239 | Sporulating parent strain | Wild-type *K. viridifaciens* | DSMZ |
| G1-G3 | Sporulating revertant | N.D. | This work |
| B1 | Revertant that recovered the ability to sporulate in the second subculture | N.D. | This work |
| B2 | Revertant with a bald colony morphology | N.D. | This work |
| B3 | Revertant with a mixed colony morphology | N.D. | This work |
| B3.1 | Revertant with a bald colony morphology | KVP1 minus, 1 Mb deletion in right arm of the chromosome | This work |
| B3.2 | Revertant with a bald colony morphology | KVP1 minus | This work |
| B3.3 | Revertant with a bald colony morphology | KVP1 minus | This work |

N.D.: Not determined

### Whole genome sequencing and analysis

For genomic DNA isolation strains were grown in Tryptic Soy Broth medium containing 10% sucrose until mid-exponential phase. Next, chromosomal DNA was isolated as described previously [23] and sequenced by BaseClear (Leiden, The Netherlands). Alignments of Illumina reads were performed using CLC Genomics Workbench 8.5.1. Raw Illumina (Hiseq2500 system) sequences of the bald strain B3.1 were imported and mapped to the reference genome of *K. viridifaciens* DSM40239 (NCBI reference sequence: NZ_MPLE00000000.1) through the “Map reads to reference” function in the NGS core tools. Mismatch cost was set to 2 and non-specific matches were handled by mapping them randomly.

### Pulsed-Field Gel Electrophoresis

For Pulsed-Field Gel Electrophoresis (PFGE), 10^6^ spores⋅ml^−1^ of *K. viridifaciens* were inoculated in 25 ml of TSBS with 0.5% glycine or LPB [25]. Cultures were grown at 30°C agitating at 200 rpm for 16 and 40 hours, respectively. Mycelial pellets were harvested by centrifugation at 4000 rpm for 15min. The preparation of plugs for PFGE was performed as previously described [21]. Plugs were made with SeaKem Gold agarose (Lonza, Switzerland), and the genomic DNA in the plugs was cut with AseI. Plugs were run using a CHEF-DR II PFGE system (Biorad, USA). For efficient separation of fragments, samples were run in two conditions: a switching time of 60-125 seconds for 20 hours, or a switching time of 2.2-75 seconds for 19 hours, both at 200 V.

### Quantitative real time PCR

Aliquots of 5 ng of DNA were used as a template in quantitative real time PCR. We used primers for both chromosomal (*atpD* and *infB*) and KVP1 (*allC*, *tetR*, *parA* and *orf1*) genes (Table 2). The PCR reactions were performed with the iTaq Universal SYBR Green Supermix Mix (Bio-Rad) using 5% DMSO, according to the manufacturer’s instructions. Reactions was performed in duplicate using a CFX96 Touch Real-Time PCR Detection System (Bio-Rad). To normalize the relative amount of DNA, the wild-type strain was used as a control, using the *atpD* gene as a reference.

**Table 2.**
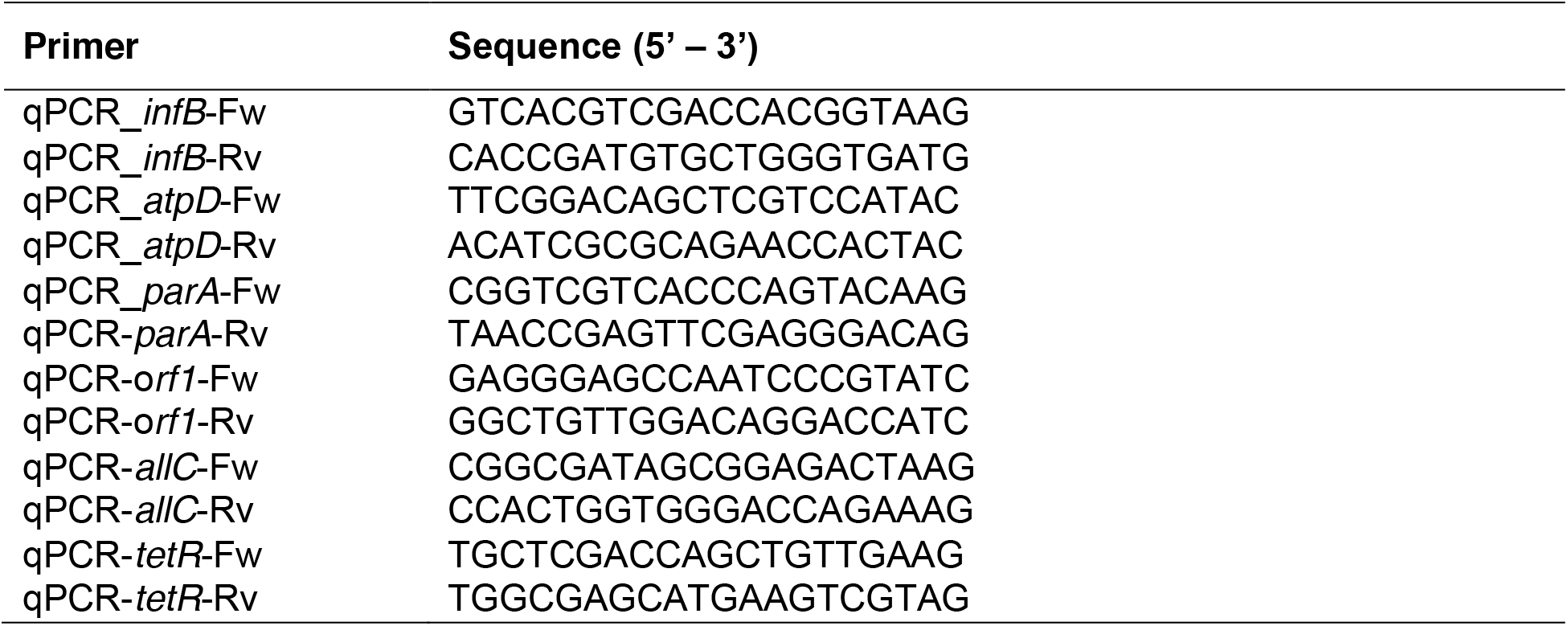
Primers used for quantitative real time PCR

## Results

### Genomic characterization of *Kitasatospora viridifaciens*

We previously sequenced *K. viridifaciens* and identified KVP1 as a novel megaplasmid [26]. Analysis of the biosynthetic gene clusters (BGCs) using antiSMASH 5.0 [27] located 11 BGCs on KVP1 and 33 clusters on the chromosome (Fig. S1). One of the BGCs showed high homology to the BGC for chlortetracycline (Fig. S2). To test experimentally whether KVP1 is indeed a plasmid, we analysed genomic DNA of the wild-type strain with Pulsed-Field Gel Electrophoresis (PFGE). In the lane containing uncut DNA, a fragment with an estimated size between 1,600,000 and 2,200,000 bp (Fig. 1A, boxed region) was evident. This fragment is consistent with a genetic element that migrates independently of the chromosomal DNA. Digestion of the DNA with AseI revealed multiple DNA fragments, including two large fragments at 1,541,168 and 1,695,864 bp (see arrowheads in Fig. 1B). By further adjusting the switching time to 2.2-75 seconds, well-separated fragments with sizes ranging from 565,000 and 945,000 bp were identified (arrowheads Fig. 1C). Combining these different PFGE runs allowed us to map the AseI fragment spectrum to the *in silico* genome assembly of *K. viridifaciens* (Fig. 1D-F). Altogether, these results confirmed that KVP1 is a megaplasmid and verified the predicted AseI sites in the chromosome.

**Figure 1.**
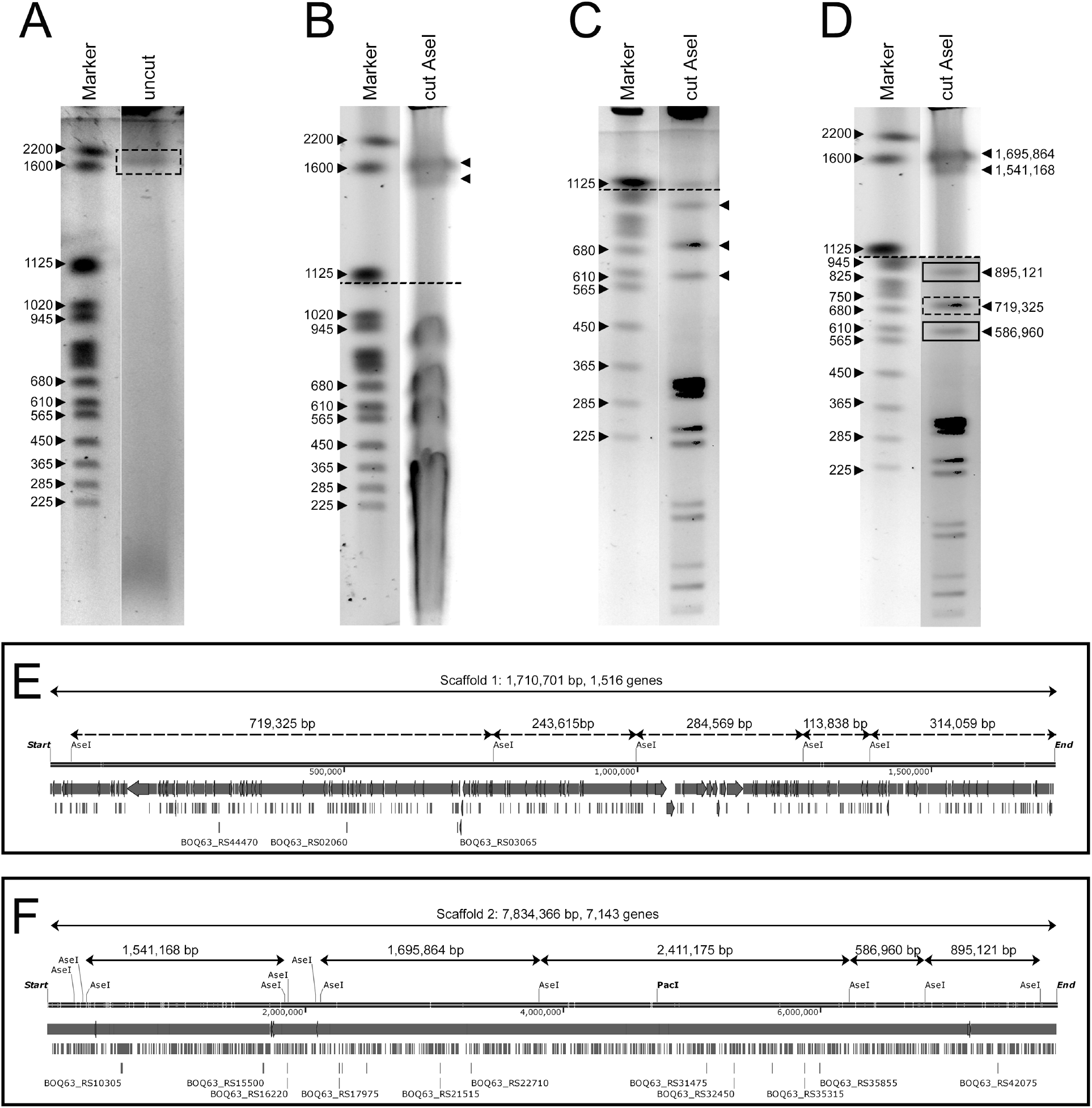
KVP1 of *K. viridifaciens* is a megaplasmid. Pulsed-Field Gel Electrophoresis of genomic DNA of *K. viridifaciens* grown in TSBS (A, C) or LPB (B) medium. DNA was separated using switching times of 60-125 seconds for 20 hours (A, B) or 2,2-75 seconds for 19 hours (C). (D) The composite gel shows fragments larger than 1,125,000 bp (derived from the gel shown in panel B), and fragments smaller than 1,020,000 bp (derived from panel C). The solid rectangles indicate fragments derived from the chromosome, while the dashed rectangles indicate fragments derived from KVP1. Predicted *in silico* maps of the KVP1 megaplasmid and chromosome of *K. viridifaciens* are shown in panels (E) and (F), respectively.

### Protoplast formation and regeneration leads to morphological diversity due to lesions and rearrangements in the chromosome and KVP1

The identification of KVP1 as a megaplasmid prompted us to analyse if it was distributed homogenously throughout the mycelium. For this, protoplasts were generated using a standard lysozyme-based protocol. Surprisingly, when protoplasts were allowed to regenerate on MYM agar plates, many colonies had developmental defects. Although the majority of colonies formed grey-pigmented spores (yellow circles Fig. 2A), a significant number of colonies was brown and failed to develop (red circles Fig. 2A). Sub-culturing of these so-called bald colonies (referred to as “B” in Fig. 2C) resulted in three morphological phenotypes, namely grey-pigmented colonies (B1), bald colonies (B2) or colonies with a variety of phenotypes (B3).

**Figure 2.**
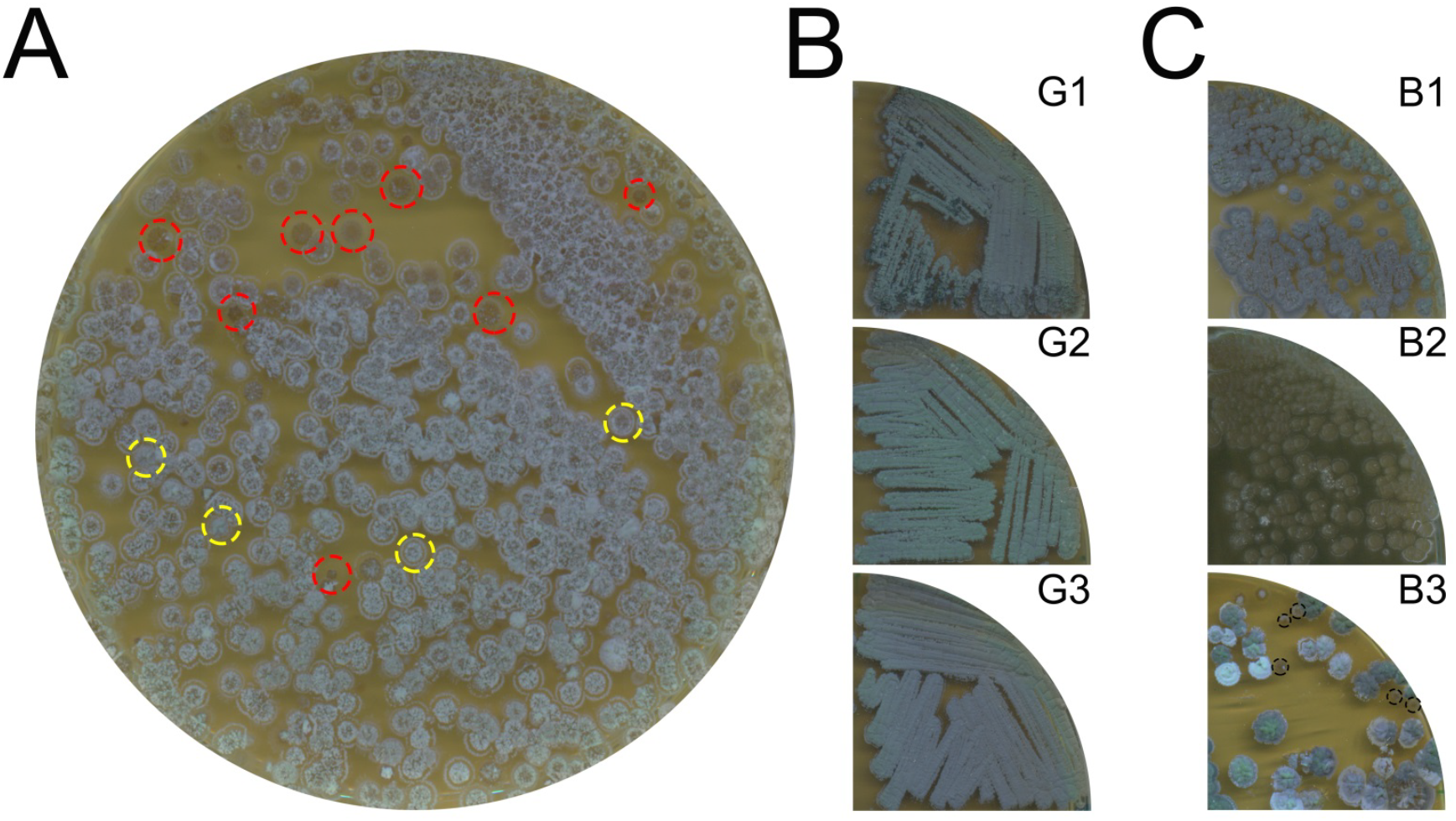
Protoplast formation and regeneration yield colonies with developmental defects. (A) Protoplasts of *K. viridifaciens* regenerated on MYM medium yields grey-pigmented colonies (yellow dotted circles) and brown colonies (red dotted circles). The grey colonies retain their morphology in subsequent subcultures (B). (C) Subculturing of the brown colonies reveals morphological heterogeneity: some colonies appear grey-pigmented (B1), while others are bald (B2) or display a variety of phenotypes (B3). Bald colonies in B3 are indicated with black dashed circles.

To rule out that the MYM medium, which lacks osmoprotectant agents, was the main cause of the morphological differences, protoplasts were also allowed to regenerate on the more commonly used R5 medium [23]. None of the colonies that arose after regeneration of protoplasts were able to differentiate on R5 medium, which is typical of *K. viridifaciens* (Fig. S3A). To analyse this further, 149 colonies were randomly selected from R5 agar plates and subsequently streaked onto MYM agar (Fig. S3B). After 7 days of growth, 77% of the colonies had a (near) wild-type morphology and produced grey-pigmented spores, while 23% of the colonies were defective in development. This demonstrates that the observed morphologically heterogeneity detected after protoplast formation and regeneration is medium-independent.

The (partial) loss of megaplasmids is known to cause morphological defects similar to those observed here [28]. To test if loss of KVP1 explains the change in phenotype, three B3-type bald colonies (encircled in Fig. 2C) were streaked onto MYM agar plates (Fig. 3A). All three colonies showed severe morphological defects and failed to produce spores even after 14 days of cultivation (Fig. 3A). Total DNA was then extracted from the three lineages (B3.1, B3.2, B3.3), and analysed for genes that served as markers for either the chromosome or for KVP1. Quantitative real-time PCR detected the chromosomal gene *infB* in the wild-type strain and the three tested lineages before the 20^th^ cycle of amplification (Fig. 3B). Conversely, the *allC* gene located on the KVP1 plasmid was only detected in the wild-type strain (Fig. 3C). Similarly, we were unable to detect other KVP1-specific genes, namely *orf1, parA* or *tetR* (Fig. 3D). These results strongly suggested the loss of KVP1 in these colonies.

**Figure 3.**
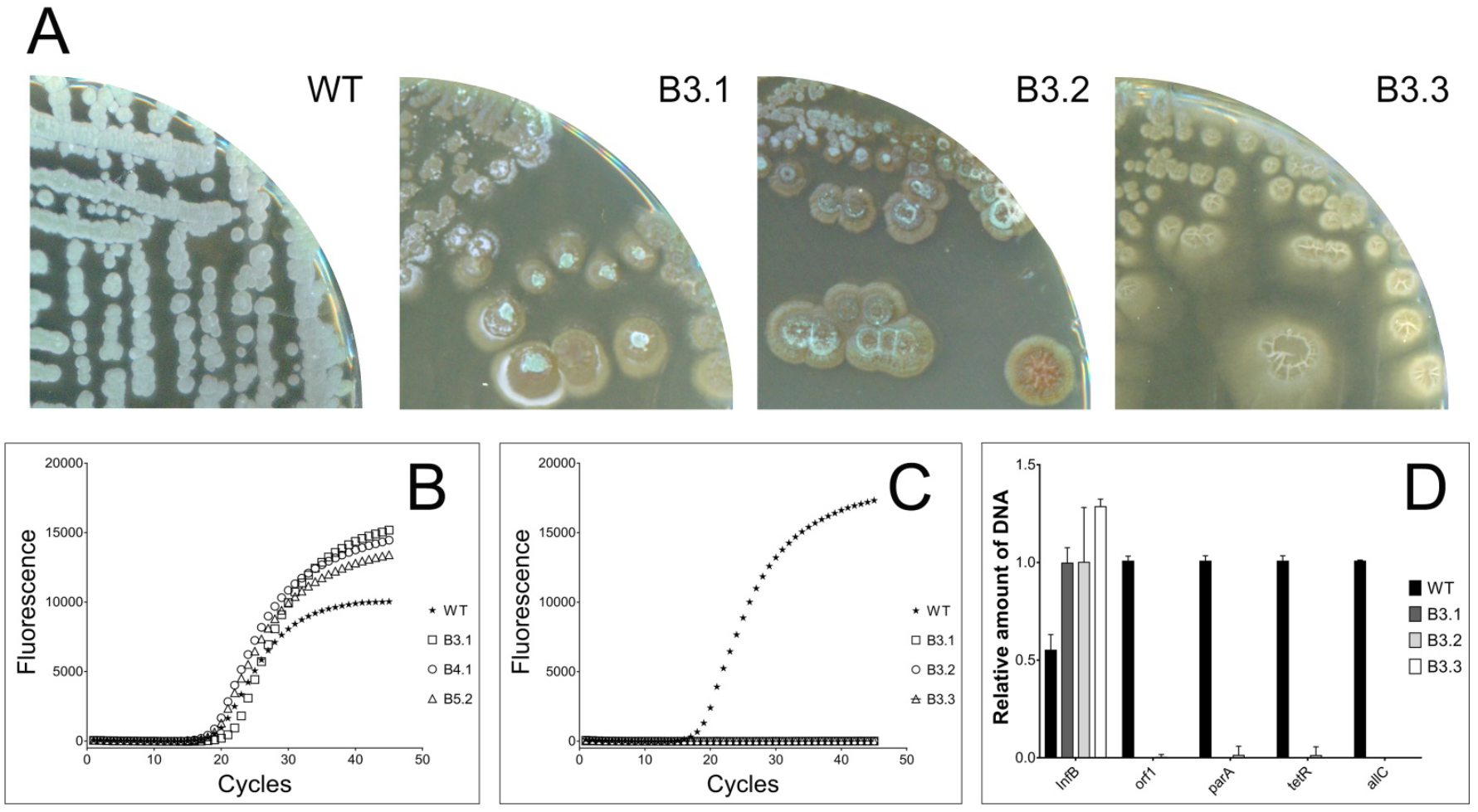
Bald colonies have lost the KVP1 megaplasmid. (A) Three independently isolated bald strains (B3.1, B3.2 and B3.3) are unable to sporulate on MYM medium, unlike the wild-type strain. Quantitative real-time PCR showed the presence of chromosomal gene *infB* in both the wild-type and bald strains (B). In contrast, the *allC* gene located on KVP1 is only present in the wild-type strain (C). (D) The relative abundance of four megaplasmid genes (*orf1*, *parA*, *tetR*, *allC*) in comparison to the abundance of the chromosomal gene *infB* suggest that KVP1 is lost in strains B3.1, B3.2 and B3.3.

To corroborate the loss of KVP1, next-generation sequencing of the total DNA of strain B3.1 was performed. While the wild-type strain showed the expected distribution of reads over genome and plasmid (3,882,545 and 1,112,768 respectively), the number of reads mapping to KVP1 was dramatically underrepresented in B3.1 (Fig. 4A, B, right panels with 5,005,998 and 143,387 reads for the chromosome and KVP1, respectively). These KVP1-mapped reads corresponded to 164,769 bp of the plasmid, mostly located on its 3’ terminal end (black box in Fig. 4A, right column). Interestingly, the number of reads mapping to the right arm of the chromosome was also dramatically decreased in B3.1 (Fig. 4B, rectangle). A more detailed analysis indicated that most chromosomal sequences between 6,261,000 bp and 7,725,700 bp (Fig. 4C, arrow) were absent in B3.1. Apparently, strain B3.1 had not only lost the majority of sequences contained on KVP1, but also a major part of its right chromosomal arm. Further investigation of the lesion sites revealed an insertion sequence (IS) immediately adjacent to the chromosomal deletion start (around 6,261,000 bp) in B3.1 (Table 3). This IS contains the BOQ63_RS37135 gene encoding the transposase likely involved in moving this element. Furthermore, close inspection of KVP1 sequences still present in strain B3.1 also identified a flanking IS element containing the BOQ63_RS06880 transposase (Table 4). Altogether, these results demonstrate that major chromosomal and megaplasmid rearrangements and DNA loss occur during protoplast formation and regeneration, which is likely mediated via transposition events.

**Table 3.** Transposition elements in the *K. viridifaciens* chromosome

| Locus | Product | Location |  |
| --- | --- | --- | --- |
|  |  | Start | End |
| BOQ63_RS09605 | IS5/IS1182 family transposase | 408,874 | 409,442 |
| BOQ63_RS10710 | transposase | 656,353 | 657,240 |
| BOQ63_RS10725 | IS256 family transposase | 659,679 | 660,950 |
| BOQ63_RS45020 | IS5/IS1182 family transposase | 1,013,156 | 1,014,156 |
| BOQ63_RS12740 | IS110 family transposase | 1,086,937 | 1,088,178 |
| BOQ63_RS12750 | IS5/IS1182 family transposase | 1,088,200 | 1,088,903 |
| BOQ63_RS15035 | IS200/IS605 family transposase | 1,569,116 | 1,569,363 |
| BOQ63_RS15720 | transposase | 1,761,237 | 1,761,554 |
| BOQ63_RS15725 | transposase | 1,761,551 | 1,762,465 |
| BOQ63_RS17065 | transposase | 2,044,875 | 2,046,032 |
| BOQ63_RS20605 | transposase | 2,847,519 | 2,848,817 |
| BOQ63_RS25160 | transposase | 3,812,365 | 3,813,189 |
| BOQ63_RS25165 | IS21 family transposase | 3,813,189 | 3,814,457 |
| BOQ63_RS25685 | transposase | 3,933,744 | 3,934,664 |
| BOQ63_RS25690 | transposase | 3,934,661 | 3,934,972 |
| BOQ63_RS26595 | IS21 family transposase | 4,116,603 | 4,117,871 |
| BOQ63_RS26600 | transposase | 4,117,871 | 4,118,695 |
| BOQ63_RS26675 | IS5/IS1182 family transposase | 4,131,168 | 4,132,061 |
| BOQ63_RS29050 | transposase | 4,605,449 | 4,606,369 |
| BOQ63_RS32225 | IS630 family transposase | 5,238,044 | 5,284,135 |
| BOQ63_RS34385 | transposase | 5,704,658 | 5,705,617 |
| **BOQ63_RS37135\*** | **IS5/IS1182 family transposase** | **6,260,623** | **6,260,976** |
| BOQ63_RS37400 | IS110 family transposase | 6,329,086 | 6,330,300 |
| BOQ63_RS40925 | transposase | 7,097,036 | 7,097,923 |
| BOQ63_RS40980 | IS5/IS1182 family transposase | 7,106,937 | 7,107,098 |
| BOQ63_RS41025 | IS481 family transposase | 7,118,421 | 7,118,739 |
| BOQ63_RS41195 | IS21 family transposase | 7,174,951 | 7,176,219 |
| BOQ63_RS41200 | transposase | 7,176,219 | 7,177,037 |
| BOQ63_RS41470 | IS110 family transposase | 7,239,487 | 7,240,701 |
| BOQ63_RS42020 | transposase | 7,367,798 | 7,369,021 |

\*Transposase found upstream chromosomal right arm deletion in bald strain B3.1

**Table 4.** Transposition elements in the right terminal region of KVP1

| Locus | Product | Location |  |
| --- | --- | --- | --- |
|  |  | Start | End |
| **BOQ63_RS06800\*** | **IS5/IS1182 family transposase** | **1,545,936** | **1,546,388** |
| BOQ63_RS06805 | IS5/IS1182 family transposase | 1,546,385 | 1,546,738 |
| BOQ63_RS06815 | transposase | 1,547,179 | 1,548,066 |
| BOQ63_RS06880 | IS5/IS1182 family transposase | 1,558,369 | 1,558,722 |
| BOQ63_RS06915 | transposase | 1,564,436 | 1,564,969 |
| BOQ63_RS06940 | IS5/IS1182 family transposase | 1,598,415 | 1,569,308 |
| BOQ63_RS07050 | IS5/IS1182 family transposase | 1,586,628 | 1,586,981 |
| BOQ63_RS07055 | IS5/IS1182 family transposase | 1,586,978 | 1,587,430 |
| BOQ63_RS07060 | IS5/IS1182 family transposase | 1,587,553 | 1,588,391 |
| BOQ63_RS07125 | IS110 family transposase | 1,600,675 | 1,600,800 |
| BOQ63_RS07220 | IS5/IS1182 family transposase | 1,621,157 | 1,622,011 |
| BOQ63_RS07225 | IS5/IS1182 family transposase | 1,622,341 | 1,623,179 |
| BOQ63_RS07245 | IS5/IS1182 family transposase | 1,624,729 | 1,625,079 |
| BOQ63_RS07255 | IS110 family transposase | 1,625,394 | 1,626,635 |
| BOQ63_RS07260 | IS5/IS1182 family transposase | 1,627,012 | 1,627,365 |
| BOQ63_RS07280 | IS5/IS1182 family transposase | 1,630,283 | 1,630,534 |
| BOQ63_RS07290 | transposase | 1,631,323 | 1,632,114 |
| BOQ63_RS07295 | transposase | 1,632,285 | 1,633,493 |
| BOQ63_RS07305 | site-specific integrase | 1,634,154 | 1,635,464 |
| BOQ63_RS07320 | DNA invertase | 1,636,936 | 1,637,601 |
| BOQ63_RS07335 | DDE transposase | 1,640,507 | 1,643,533 |
| BOQ63_RS07345 | integrase | 1,645,074 | 1,646,333 |
| BOQ63_RS07380 | integrase | 1,652,931 | 1,654,022 |
| BOQ63_RS07400 | resolvase | 1,656,203 | 1,656,996 |
| BOQ63_RS07425 | transposase | 1,658,975 | 1,659,889 |
| BOQ63_RS07430 | DDE transposase | 1,660,210 | 1,661,212 |
| BOQ63_RS07440 | transposase | 1,661,565 | 1,662,452 |
| BOQ63_RS07445 | IS30 family transposase | 1,662,486 | 1,662,824 |
| BOQ63_RS07475 | IS21 family transposase | 1,665,953 | 1,667,221 |
| BOQ63_RS07480 | IS21 family transposase | 1,667,436 | 1,668,701 |
| BOQ63_RS07485 | transposase | 1,668,701 | 1,669,546 |
| BOQ63_RS07550 | transposase | 1,689,112 | 1,689,999 |
| BOQ63_RS07585 | transposase | 1,695,301 | 1,695,435 |
| BOQ63_RS07615 | IS5/IS1182 family transposase | 1,699,532 | 1,699,984 |
| BOQ63_RS07620 | IS5/IS1182 family transposase | 1,699,981 | 1,700,334 |
| BOQ63_RS07635 | transposase | 1,704,820 | 1,706,307 |
| BOQ63_RS07640 | TnsA-like heteromeric transposase | 1,706,304 | 1,706,978 |

\*Transposase found upstream KVP1 right terminus present in bald strain B3.1

**Figure 4.**
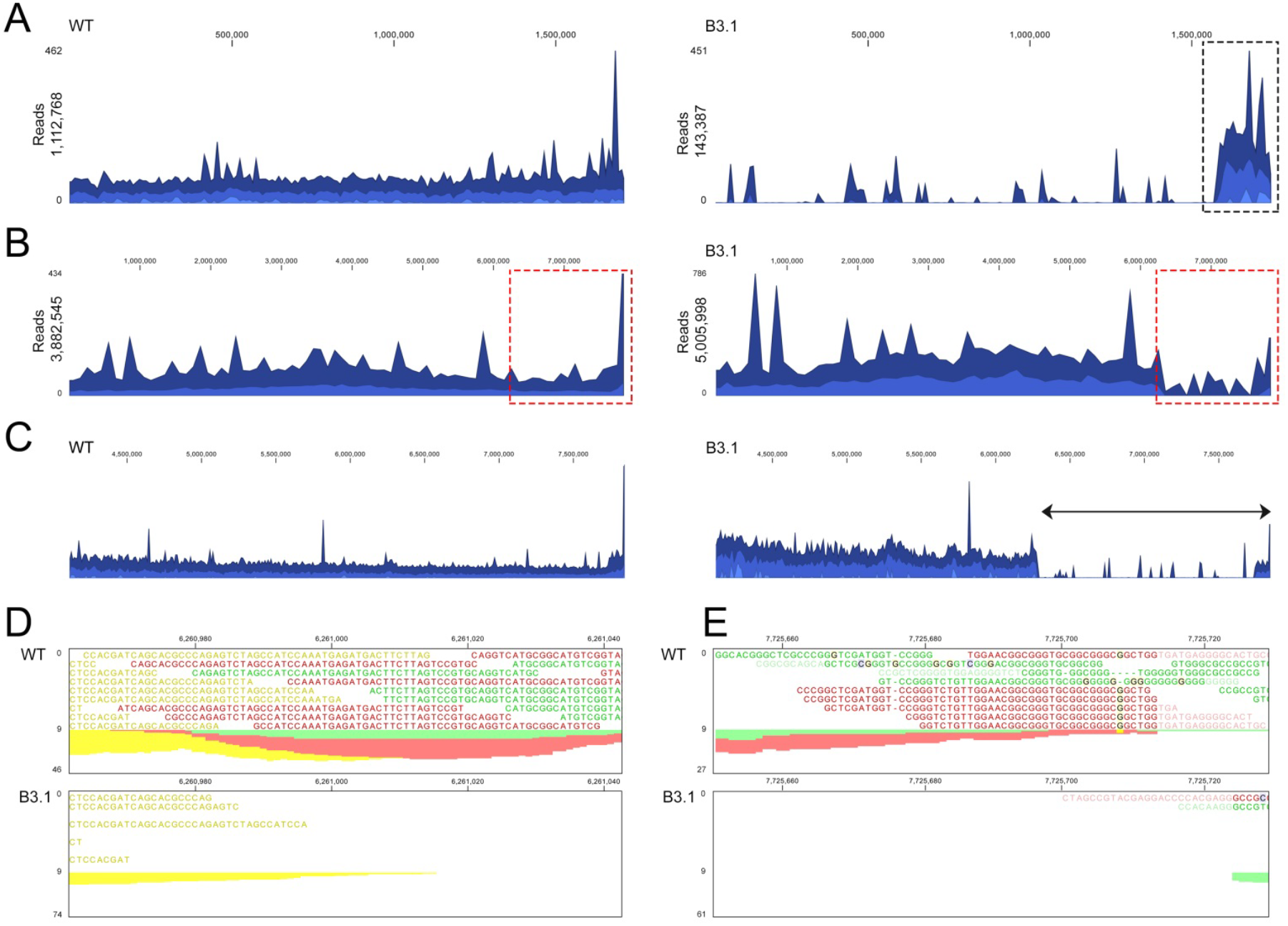
Whole genome sequencing reveals major chromosomal and megaplasmid lesions in strain B3.1. Alignment of Illumina reads of the wild type (left) and B3.1 strain (right) against KVP1 (A) and the chromosome (B). Please note the high coverage of KVP1 sequences (panel A) detected in the wild type (1,112,768 reads) compared to those of strain B3.1 (143,387 reads). Similarly, a high coverage in the right arm of the chromosome (dashed red rectangle) is observed for the wild-type (B, left panel) in comparison to strain B3.1 (B, right panel). (C-E) A more detailed characterization reveals that all reads between 6,261,000 and 7,725,700 are absent in the right arm of the chromosome of strain B3.1

## Discussion

Actinobacterial genomes readily undergo rearrangements, whereby more than 1 Mb of genomic DNA can be lost [18, 21, 29]. Here, we provide evidence that protoplast formation and regeneration in *K. viridifaciens* can lead to profound genomic rearrangements in the chromosome as well as loss of (large parts of) the megaplasmid KVP1. Given that these genomic rearrangements translate into major phenotypic variations, caution should be taken when using protoplasts for creating mutants, in particular when using strains that carry natural plasmids.

Filamentous Actinobacteria grow by tip extension and develop multinucleated mycelia. Little is known on how these bacteria regulate the abundance and spatial distribution of chromosomes and extrachromosomal plasmids within the mycelium. Maintaining large plasmids such as KVP1 is costly, given that such elements can comprise about a fifth of the entire genome. Megaplasmids are often reservoirs for biosynthetic gene clusters, and their interactions with the chromosome have been suggested to be a driving force for horizontal gene transfer [13]. Loss of such plasmids not only affects morphological development but may also influence the production of specialized metabolites whose gene clusters are contained on the chromosome. In *Streptomyces hygroscopicus* elimination of pSHJG1 increased the production of validamycin A [30], while holomycin yield was boosted when pSCL4 was lost in *Streptomyces clavuligerus* [28, 31]. Whether the loss of KVP1 has a similar effect on production of specialized metabolites in *K. viridifaciens* remains to be elucidated.

We observed that close to one fourth (23%) of the colonies derived from regenerated protoplasts of *K. viridifaciens* were defective in aerial growth and sporulation. Similar morphological defects have been described for *S. clavuligerus*, *Streptomyces lividans* and *Streptomyces coelicolor* upon loss of their plasmids [28, 32]. While the loss of KVP1 in *K. viridifaciens* may explain the arrest in morphological development, we here show that such plasmid-lacking derivatives can also carry other profound lesions in the chromosome, which could equally well contribute to this phenotype. By sequencing one revertant that had lost KVP1 we found that this strain had also lost approximately 1.5 Mb of the right arm of the chromosome. Such genetic instability is typical of streptomycetes, and can affect morphological differentiation, but also phenotypic traits associated with natural products, such as pigmentation, antibiotic biosynthesis and antibiotic resistance [20, 21]. Transposable elements were suggested as the principal cause of genetic instability [33]. The loss of KVP1 and the chromosomal lesions in the right arm could be the consequence of replicative transposition between the chromosome and the megaplasmid (Fig. 5). Notably, while most KVP1-located sequences were absent in the sequenced strain, including those required for autonomous replication of this megaplasmid, we identified a high coverage of sequences originally located at the 3’ end of KVP1. This could be explained by an exchange of the 3’ end of KVP1 with the right chromosomal arm. A replicative transposition event between two linear replicons often results in the loss of chromosomal terminal regions and recombination of transposable elements [19]. The presence of IS elements located at the chromosomal and plasmid termini could provide the basis for homologous recombination between these DNA molecules. It has been previously shown that homologous copies of IS elements could serve as substrates for the recombination machinery, creating chromosomal rearrangements in the genomes of *Lactococcus lactis* and *Escherichia coli* [34, 35]. In the case of colony B3.1 that had arisen from protoplast regeneration, our results suggest that a possible cause of the genomic rearrangement was the replicative transposition of an IS, exerted by the BOQ63_RS37135 transposase (black arrow in Fig. 5B). Following replicative transposition, a double-stranded break occurs at the site of transposon excision. This break is repaired by recombination with homologous genes located on IS elements present in the right arm of KVP1 (shown as a grey arrow in the KVP1 right arm), which are abundantly present. This recombination might force the interchange of terminal arms. This hypothetical model would explain the genome size reduction of B3.1 (6,787,546 bp).

**Figure 5.**
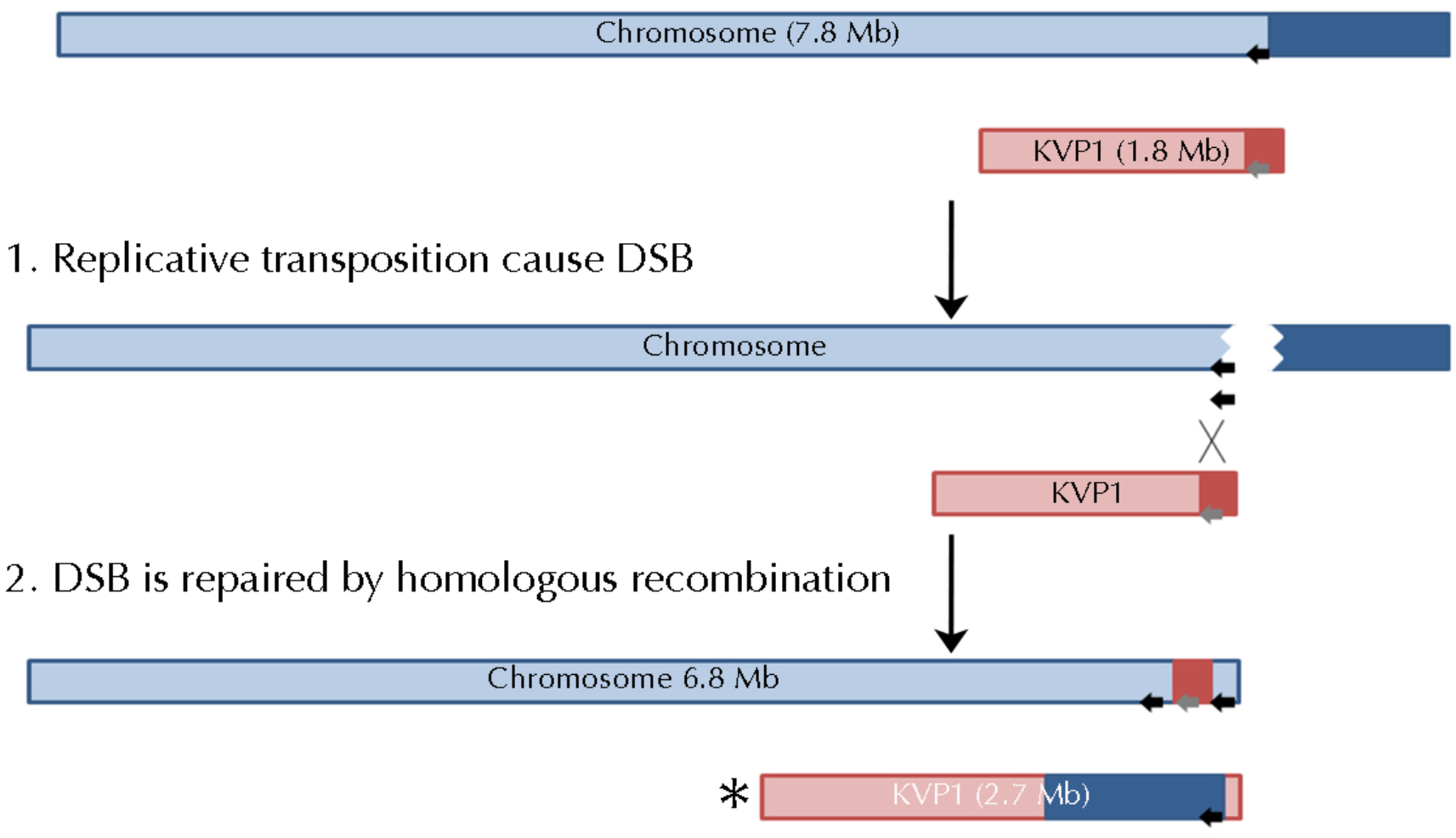
Proposed model for the genomic rearrangements and lesions identified in strain B3.1. Replicative transposition of a transposable element (black arrow) located on the chromosome creates a double-stranded break (DSB). Subsequent repair of the DSB by homologous recombination between the chromosome and KVP1 plasmid (carrying numerous transposable elements), leads to an exchange of the right arms of both replicons and changes in their size. Please note that the larger KVP1 megaplasmid variant is lost in strain B3.1.

The frequency of aberrant phenotypes after protoplast regeneration is higher than the phenotypic heterogeneity obtained after outgrowth of spores, which typically is in the order of 1% [18, 21, 29, 33, 36, 37]. An explanation for the high frequency of aberrant mutants in colonies arising after protoplasting may relate to the activation of transposable elements contained in the terminal regions of the chromosome and/or the KVP1 plasmid. The activation of transposases are typically stimulated by stressful conditions, such as radiation, oxidative stress, temperature or inhibitory concentrations of metals and antibiotics [38]. It was recently demonstrated that elevated levels of osmolytes induces hyperosmotic stress [25, 39], which are conditions that are also used during preparation of protoplasts. This stress could also stimulate transposition events and consequent chromosomal rearrangements. Consistent with this idea is that other cell wall-deficient cells, called L-forms, which have likewise been exposed to osmotic stress conditions, carry chromosomal lesions. In this context it is interesting to note that in three independent L-form lineages of *K. viridifaciens* lesions in the right chromosomal arms were found in addition to loss of KVP1 [25]. These three strains retained a similar region of KVP1 in their genomes, with a size of 164,773 bp for *alpha* and M1, and 164,642 bp for M2. These are very similar to the KVP1-sequences remaining (164,769 bp) in the bald protoplast regenerant B3.1 in terms of length and content.

Chromosomal rearrangements are often detrimental for the fitness of a unicellular organism. However, it was recently shown that in *Streptomyces* chromosomal rearrangements may increase the diversity and production of specialized metabolites, including antibiotics [21]. A division of labour strategy would allow a colony to have a mixture of mutant and wild-type chromosomes, where the mutant cells are virtually sterile and become specialized in the production of antibiotics, while the cells containing wild-type chromosomes are efficient spore producers [21]. Thus, while some genetic variation may naturally exist within the mycelium, we expect that exposure to high levels of osmolytes, associated with growth and subsequent protoplast formation, generates stress and dramatically increases chromosomal changes. This study provides a starting point to further characterize these changes and to investigate their consequences, which may lead to exciting new insights into the biology of these prolific antibiotic producers.

## Author statements

K.R. and Z.Z. collected the data and aided in data analysis. K.R. and D.C. designed the experiments. D.E.R., G.P.W. and D.C. supervised the research. K.R. and D.C. wrote the paper with input from all co-authors.

---

**Figure S1.**
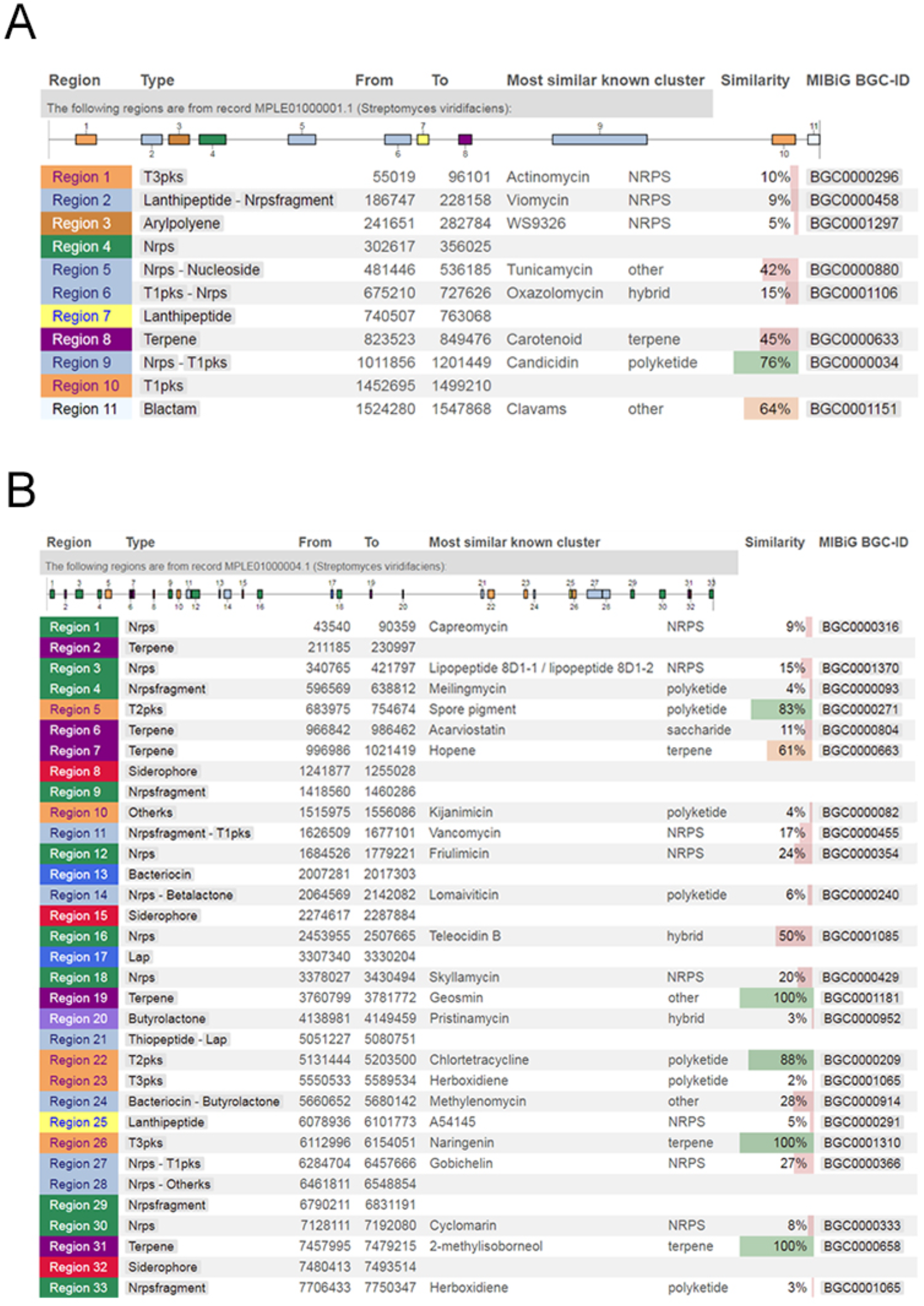
AntiSMASH 5.0 output revealing the biosynthetic gene clusters contained on the megaplasmid (top) and chromosome (bottom) of *K.viridifaciens*. The biosynthetic gene clusters are numbered according to their localization on the replicon.

**Figure S2.**
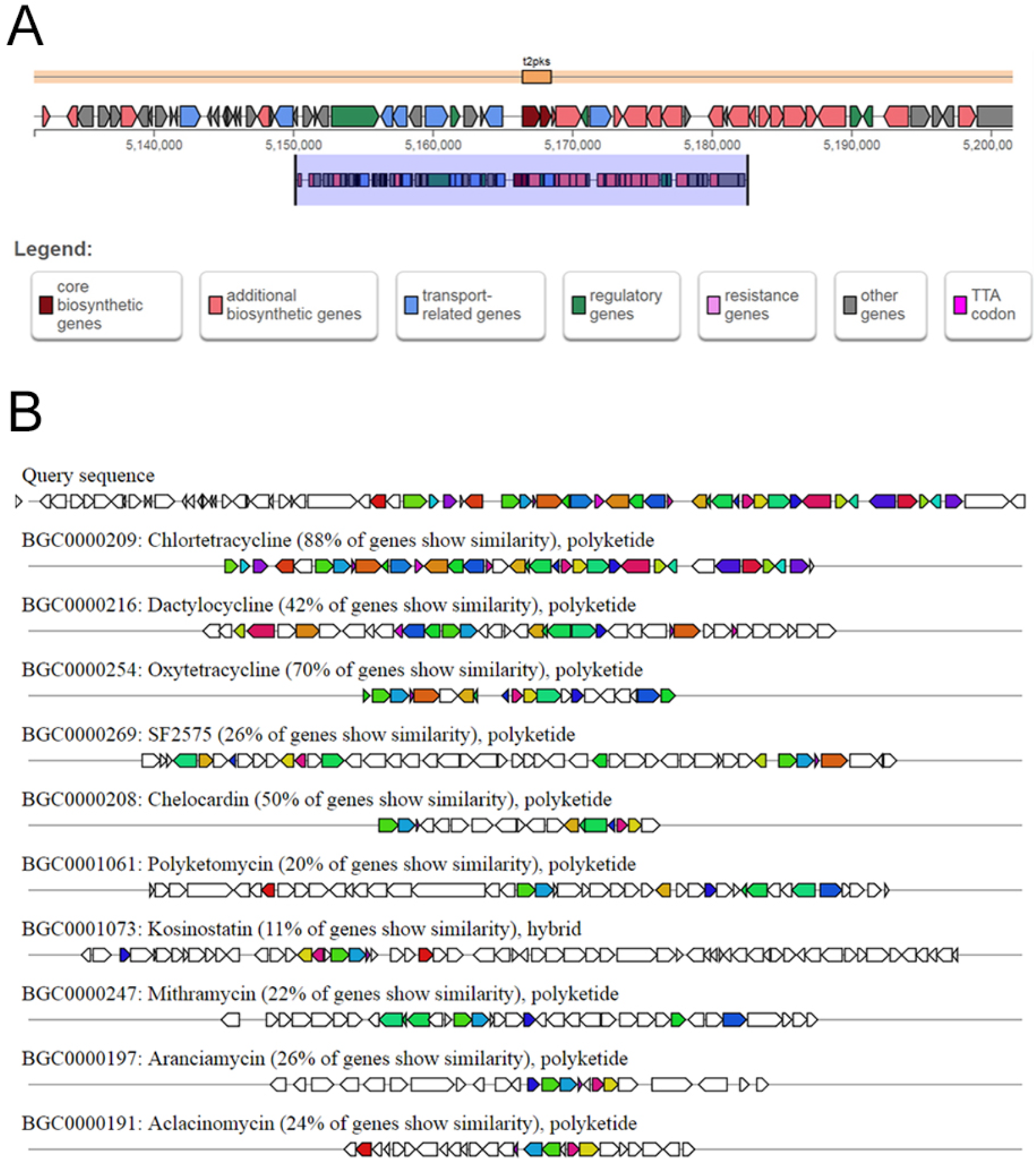
AntiSMASH 5.0 homology search of the *K. viridifaciens* tetracycline biosynthetic gene cluster. (A) Localization of the putative chlorotetracycline BGC in the chromosome of *K. viridifaciens*. (B) Comparison of the *K. viridifaciens* tetracyclin biosynthetic gene cluster with known tetracycline gene clusters from *Streptomyces aureofaciens* (BGC0000209), *Dactylosporangium sp.* SC14051 (BGC0000216), *Streptomyces rimosus* (BGC0000254), *Streptomyces sp*. SF2575 (BGC0000269), *Amycolatopsis sulphurea* (BGC0000208), *Streptomyces diastatochromogenes* (BGC0001061), *Micromonospora sp*. TP-A0468 (BGC0001073), *Streptomyces argillaceus* (BGC0000247), *Streptomyces echinatus* (BGC0000197) and *Streptomyces galilaeus* (BGC0000191).

**Figure S3.**
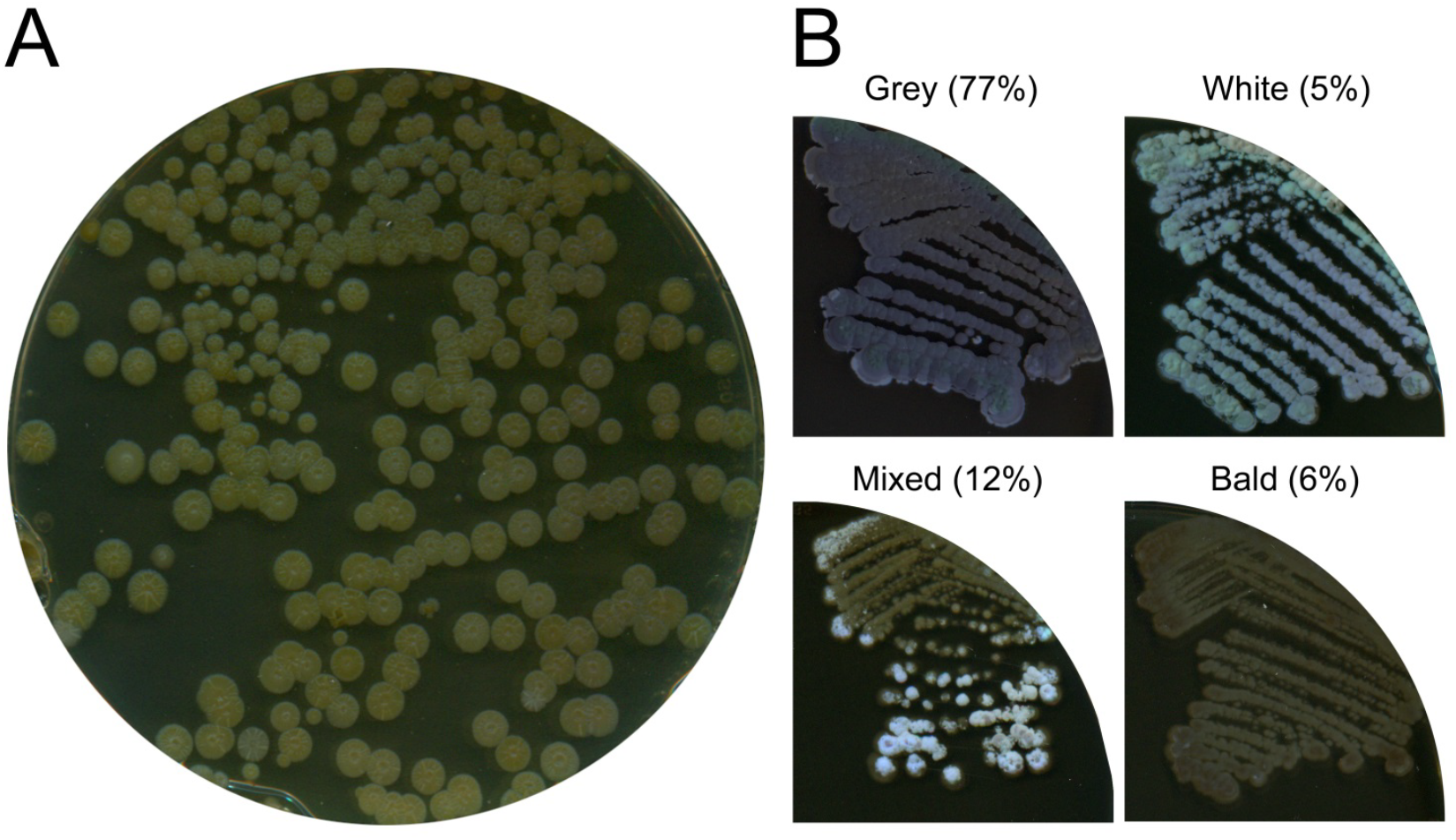
Protoplast regeneration generates morphological diversity in osmotically balanced medium. (A) Protoplasts regenerated on R5 medium yielded colonies that are unable to sporulate due to the high sucrose levels. (B) Subculturing of 149 randomly-picked colonies on MYM medium revealed dramatic developmental defects in 23% of the colonies. Whereas 77% of the colonies were able to form grey-pigmented sporulating colonies (similar to the wild-type), 5% of the colonies were white, 6% were bald, while 12% had a mixed appearance.

